# The interaction of enteric bacterial effectors with the host engulfment pathway control innate immune responses

**DOI:** 10.1101/2021.06.07.447447

**Authors:** Ibrahim M Sayed, Stella-Rita Ibeawuchi, Dominique Lie, Mahitha Shree Anandachar, Rama Pranadinata, Manuela Raffatellu, Soumita Das

**Author notes:** **Address correspondence to**: Soumita Das, Department of Pathology, University of California, San Diego, 9500 Gilman Drive, MC 0644, George Palade laboratory, Office Rm 256, San Diego, CA, 92093-0644, USA, E mail. Another Affiliation: Department of Medical Microbiology and Immunology, Faculty of Medicine, Assiut University, Egypt. **Author Contributions:** Study concept and design: IMS, SRI, DL, MSA, RP, MR, SD Acquisition of data: IMS, SRI, DL, MSA, RP Analysis and interpretation of data; IMS, SRI, DL, MSA. Drafting of the manuscript: IMS, SD Critical revision of the manuscript for important intellectual content: IMS, SRI, MSA, RP, MR, SD Statistical analysis: IMS, SD Obtained funding: SD Study supervision: SD.

## Abstract

**Background:** Host engulfment protein ELMO1 generates intestinal inflammation following internalization of enteric bacteria. In *Shigella*, bacterial effector IpgB1 interacts with ELMO1 and promotes bacterial invasion. IpgB1 belongs to the WxxxE effector family, a motif found in several effector of enteric pathogens. Here, we have studied the role of WxxxE effectors, with emphasis on *Salmonella* SifA and whether it interacts with ELMO1 to regulate inflammation.

**Methodology:** In-silico-analysis of WxxxE effectors was performed using BLAST search and Clustal W program. The interaction of ELMO1 with SifA was assessed by GST pulldown assay and co-immunoprecipitation. ELMO1 knock-out mice, and ELMO1-depleted murine macrophage J774 cell lines were challenged with WT and *SifA* mutant *Salmonella*. Bacterial effectors containing the WxxxE motif were transfected in WT and ELMO1-depleted J774 cells to assess the inflammatory cytokines.

**Results:** ELMO1 generates differential pro-inflammatory cytokines between pathogenic and non-pathogenic bacteria. WxxxE motif is present in pathogens and in the TIR domain of host proteins. The C-terminal part of ELMO1 interacts with SifA where WxxxE motif is important for interaction. ELMO1-SifA interaction affects the bacterial colonization, dissemination, and inflammatory cytokines *in vivo*. Moreover, ELMO1-SifA interaction increases TNF-α and IL-6 production from the macrophage cell line and is associated with enhanced Rac1 activity. ELMO1 also interacts with WxxxE effectors IpgB1, IpgB2, and Map, and induces inflammation after challenge with microbe or microbial ligand.

**Conclusion:** ELMO1 generates a differential response through interaction with the WxxxE motif which is absent in commensals. ELMO1-WxxxE interaction plays a role in bacterial pathogenesis and induction of inflammatory response.

**Highlights:**

1. ELMO1 generates a differential immune response between enteric pathogens and commensals.
2. Enteric bacterial effectors containing WxxxE signature motif interact with ELMO1.
3. The WxxxE effector of *Salmonella* SifA interacts with the C-terminal part of ELMO1.
4. ELMO1-SifA interaction increases the inflammatory response *in vivo* and *in vitro*.

## Introduction

Host defense detects the presence of harmful bacteria to initiate a protective response. The host immune cells recognize the pathogen-associated molecular patterns (PAMPs) of bacteria through their pattern recognition receptors (PRRs) (Akira et al., 2006; Ansaldo et al., 2021; Evans, 2009; Liwinski et al., 2020; Perez-Lopez et al., 2016; Sassone-Corsi and Raffatellu, 2015). Although lipopolysaccharide (LPS) is a key cell wall component of both pathogenic and commensal Gram-negative bacteria, host defenses exhibit a differential immune response against bacteria which affect the ability to cause disease (Sansonetti, 2011; Stuart et al., 2013). The mechanisms by which host sensors of microbe can differentiate between pathogens and commensals are not completely recognized.

Previously, we showed that the host engulfment protein called EnguLfment and cell MOtility protein 1 (ELMO1) plays a crucial role in the internalization of enteric pathogens, regulation of autophagy induction, and bacterial clearance during enteric infection (Das et al., 2015; Sarkar et al., 2017). ELMO1 is a cytosolic protein that interacts with another PRR called Brain Angiogenesis Inhibitor 1 (BAI1) which binds with the oligosaccharide core of the LPS of Gram-negative bacteria (Das et al., 2011; Das et al., 2014). ELMO1 also interacts with the cytosolic protein Dock180 and together act as a guanine nucleotide exchange factor for the small Rho GTPase Ras-related C3 botulinum toxin substrate 1 (Rac1), leading to actin cytoskeleton reorganization and bacterial engulfment (Das et al., 2011; Das et al., 2015). The polymorphism of ELMO1 is involved in several inflammatory diseases such as inflammatory bowel disease, rheumatoid arthritis, kidney disease, and diabetic nephropathy (Parmar et al., 2012; Pezzolesi et al., 2009; Whitaker et al., 2015). Recently, we showed that ELMO1 expression is elevated in the colonic epithelium of IBD patients, where higher expression is positively correlated with the elevated expression of pro-inflammatory cytokines, MCP-1, and TNF-α (Sayed et al., 2020). However, whether ELMO1 differentially initiates the immune response after sensing pathogens and/or commensals remains to be determined.

To survive and proliferate inside the host, pathogens utilize a variety of secretion systems (types I–VI) that target host proteins to hijack host defense mechanisms (Alto et al., 2006; Cornelis and Van Gijsegem, 2000; Galán and Collmer, 1999). Some bacterial effectors target the host cytoskeleton G protein signaling cascades of the Rho family of GTPases resulting in induction of cytoskeletal rearrangements and facilitating the bacterial entry (Hardt et al., 1998; Shao et al., 2002). Previously it has been shown that ELMO1 interacts with IpgB1, a WxxxE effector of *Shigella* and controls internalization in epithelial cells (Handa et al., 2007). Similar to IpgB1 and IpgB2 of *Shigella*, other bacterial effectors such as Map from enteropathogenic *Escherichia coli* (EPEC), and SifA from *Salmonella*, bypass the endogenous RhoGTPases to directly activate downstream signaling responses (Alto et al., 2006). These effectors have a conserved sequence similarity in their C-terminal targeting sequences entitled “WxxxE motif” (Alto et al., 2006). The WxxxE motif is reported in 24 effector proteins present in enteric pathogens and they all can utilize a similar molecular mechanism (Alto et al., 2006). However, whether the WxxxE motif is unique to pathogens, and if so whether it could promote an immune response mediated by ELMO1 that discriminates between pathogens and commensals, is not known.

Here we investigated whether ELMO1 interacts with other WxxxE effectors and how the effector-ELMO1 interaction controls inflammatory responses. Our in-silico analysis identified WxxxE signature motif to be predominantly present in enteric pathogens, plant pathogens, and in the Toll/interleukin-1 receptor (TIR) homology domain of both host and pathogens but absent in commensals. Using *Salmonella* as a model for enteric pathogens, we found that the WxxxE motif is important to maintain the interaction between the *Salmonella* SifA effector and ELMO1. ELMO1-KO mice and ELMO1-depleted macrophages were used to study the impact of ELMO1-SifA interaction on bacterial colonization, dissemination, and inflammatory response. ELMO1 also interacts with WxxxE effectors from *Shigella* and *E. coli* and the interaction of ELMO1 with WxxxE effectors induces the production of pro-inflammatory cytokines.

## RESULTS

### ELMO1 generates a differential immune response between pathogenic and non-pathogenic bacteria

We have previously shown that ELMO1 is involved in the internalization of *Salmonella* ((Das et al., 2015)). To understand the role of ELMO1 in the internalization of pathogenic and non-pathogenic bacteria, we infected control and ELMO1 shRNA J774 macrophages with the pathogen *Salmonella* and with two non-pathogenic *E. coli* strains (K12 and DH5α). Gentamicin protection assay showed that the number of internalized bacteria was low in control shRNA cells after infection with the non-pathogenic *E. coli* strains compared to *Salmonella* **(Figure 1A)**. Bacterial internalization was lower in ELMO1 shRNA cells compared to control cells irrespective of pathogenic or non-pathogenic bacteria. The percentage internalization **(Figure 1B**) showed ELMO1-dependent bacterial internalization in all three cases, with a significant ∼50% reduction of internalization in ELMO1 shRNA cells compared to control cells.

**Figure 1:**
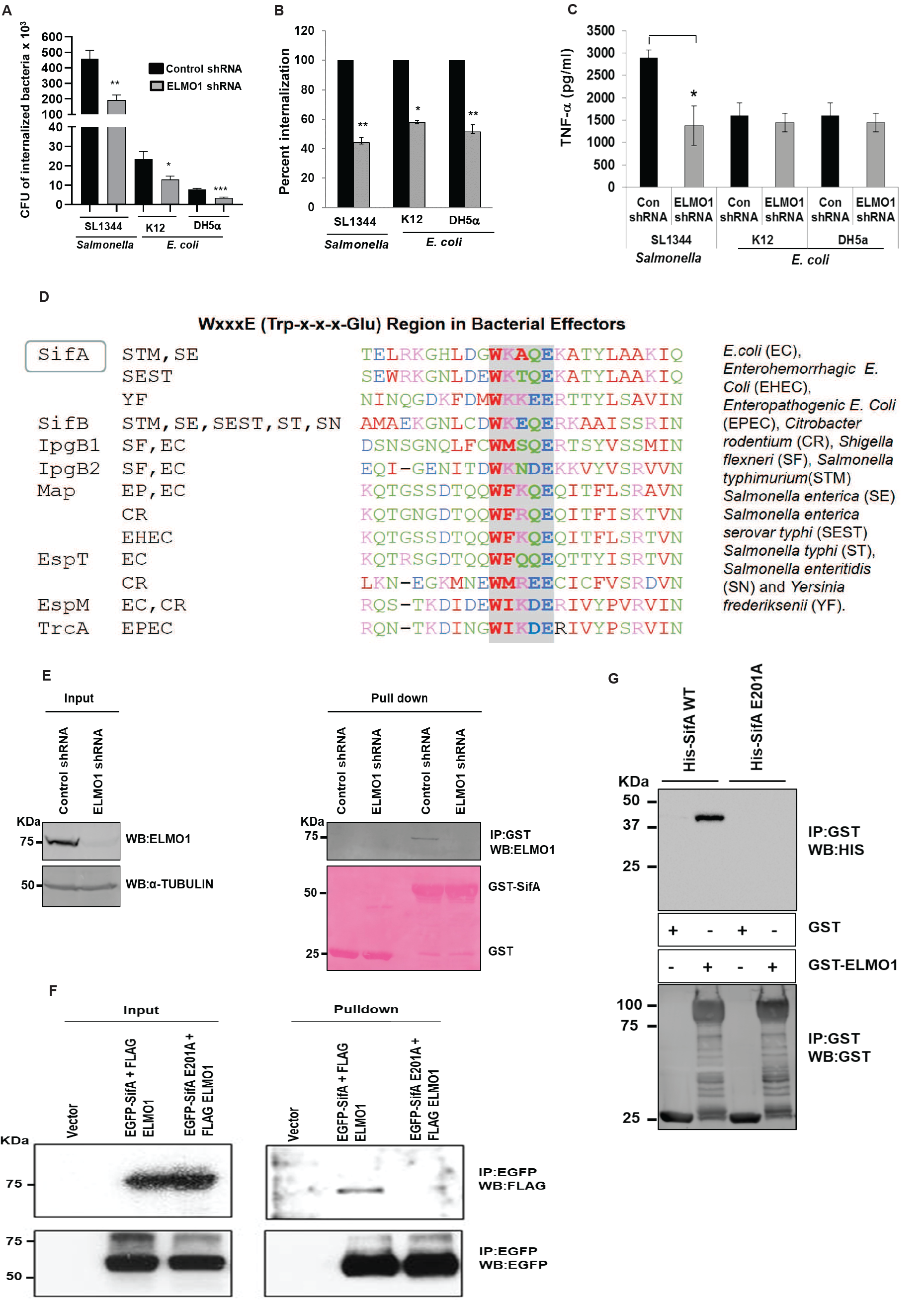
ELMO1 generates a differential immune response between pathogenic and non-pathogenic bacteria. (A) Control and ELMO1 shRNA J774 cells were challenged with *Salmonella* enterica serovar Typhimurium (SL1344) as a source of pathogenic bacteria, and *E. coli* strain K12 and *E. coli* DH5α as a source of non-pathogenic bacteria for 1 h at 37°C. Bacterial internalization was measured using gentamicin protection assay as described in Materials and Methods. (B) The percentage of bacterial internalization was compared between control and ELMO1 shRNA J774 cells, bacterial internalization for control cells was taken as 100% as performed in A. (C) The level of TNF-α was measured in control and ELMO1 shRNA J774 cells challenged with SL1344, *E. coli* K12, and *E. coli* DH5α after 3 h. Data in (A), (B), and (C) represent the mean ± SEM of three independent experiments. *, **, *** means *p* ≤ 0.05, ≤ 0.01, and ≤ 0.001, respectively, as assessed by unpaired two-tailed Student t-test. (D) BLAST-Protein search using the amino acid sequence of *Shigella* IpgB1 with all other bacteria identified sequence similarities with a signature motif (WxxxE or Trp-x-x-x-Glu) present in bacterial effectors from enteric pathogens but absent in commensals. (E) GST pulldown was performed with control GST and GST-SifA with the lysates from control and ELMO1 shRNA J774 cells. The input was shown on the left side and immunoblotted with anti-ELMO1 antibody. α -Tubulin was used as a loading control. (F) HEK 293 cells were transfected with either the vector control or with the EGFP-SifA (with WxxxE signature motif)+FLAG-ELMO1 or with the EGFP-SifAE201A (WxxxE mutant motif)+FLAG-ELMO1. The lysates from each condition were used either as input (left) or used for EGFP pulldown, followed by immunoblotting with anti-FLAG antibody to know the level of FLAG-ELMO1. Equal loading is confirmed by the EGFP antibody (lower panel). (G) GST pulldown with either the GST alone or with GST-ELMO1 was incubated with His-SifA WT (with WxxxE signature motif) and His-SifA E201A (WxxxE mutant motif). The pulldown samples were immunoblotted with anti-His antibody (upper panel). The equal loading of beads is confirmed by anti-GST antibody (lower panel).

To assess whether ELMO1 generates a differential immune response between pathogenic and non-pathogenic bacteria, we measured the level of pro-inflammatory cytokine TNF-α released from control and ELMO1 shRNA cells. TNF-α levels were significantly lower in ELMO1 shRNA cells compared to control cells infected with *Salmonella*. However, the level of TNF-α was comparable in control and ELMO1 shRNA cells infected with non-pathogenic *E. coli* **(Figure 1C)**. This finding suggests that this ELMO1-dependent cytokine response is likely triggered by a virulence factor from *Salmonella*. Previously, we have shown that the TNF-α response in macrophages after interaction with microbial ligands (LPS and lipoteichoic acid) does not depend on ELMO1 (Das et al., 2015). While searching for bacterial factors potentially involved in the ELMO1-dependent differential cytokine responses, we found a previous report where ELMO1 interacts with the *Shigella* effector IpgB1 (Handa et al., 2007). Of note, IpgB1 belongs to the WxxxE effector groups and shares the signature motif WxxxE (Tryptophan-xxx-Glutamate) found only in enteric pathogens (Alto et al., 2006). A BLAST database search identified the conserved signature WxxxE motif as widely distributed among enteric pathogens **(Figure 1D, Supplement Figure 1A)**, non-enteric pathogens including *Acinetobacter* and *Klebsiella* (**Supplement Figure 1B)**, and plant pathogens (**Supplement Figure 1C)**, but not among commensals. Interestingly, the WxxxE motif is also present in the TIR domain of a subset of host TLRs (TLR 1, 4, 6, 7, 8, 9, 10), and TLR1, 4, 6, 10 have 2 WxxxE motifs (**Supplement Figure 1D**).

Based on the bioinformatics analysis that identified the WxxxE motif in the *Salmonella* effector SifA (**Figure 1D**), we examined whether *Salmonella* SifA interacts with endogenous ELMO1 in macrophages **(Figure 1E)**. GST pulldown assay showed ELMO1 in J774 control shRNA cells bound to GST-SifA and the specificity was further confirmed when J774 ELMO1 shRNA cells failed to bind the GST-SifA. To further test the impact of WxxxE signature motif, we performed immunoprecipitation with FLAG-tagged-ELMO1 in cells co-transfected with either EGFP-SifA wild type or EGFP-SifA-E201A mutant. We chose the mutation of the glutamine (E) but not the tryptophan (W) residue because a previous study showed that a mutation in W of the WxxxE motif affects the stability and/or tertiary structure of SifA and impedes its interaction with SKIP (Diacovich et al., 2009). Mutation of the WxxxE_201_ motif abolished the interaction between SifA and ELMO1 when we performed the immunopulldown (IP) with EGFP-conjugated beads **(Figure 1F)**. We further confirmed the specificity of the interaction by GST pulldown with purified His-tagged SifA WT or E201A mutant and GST ELMO1 **(Figure 1G)**. The results indicate the WxxxE motif is important for the interaction of SifA with ELMO1.

### ELMO1-SifA interaction affects bacterial dissemination and inflammatory responses *in vivo*

By using ELMO1-knockout (KO) mouse, we have previously shown that ELMO1 promotes *Salmonella* dissemination and intestinal inflammation (Das et al., 2015). To assess the relevance of ELMO1 and SifA interaction on bacterial gut colonization, dissemination, and inflammatory responses *in vivo*, WT and global ELMO1 KO mice were infected with *Salmonella enterica* serovar Typhimurium wild-type strain SL1344 (WT *SL*) or with an isogenic *sifA* mutant with a dose of 5×10^7^ CFU/mouse for 5 days by oral gavage. Samples from the ileum, cecum, spleen, and liver were collected to measure bacterial burden and the level of inflammatory cytokines as shown in the schematics in **Figure 2A**.

**Figure 2.**
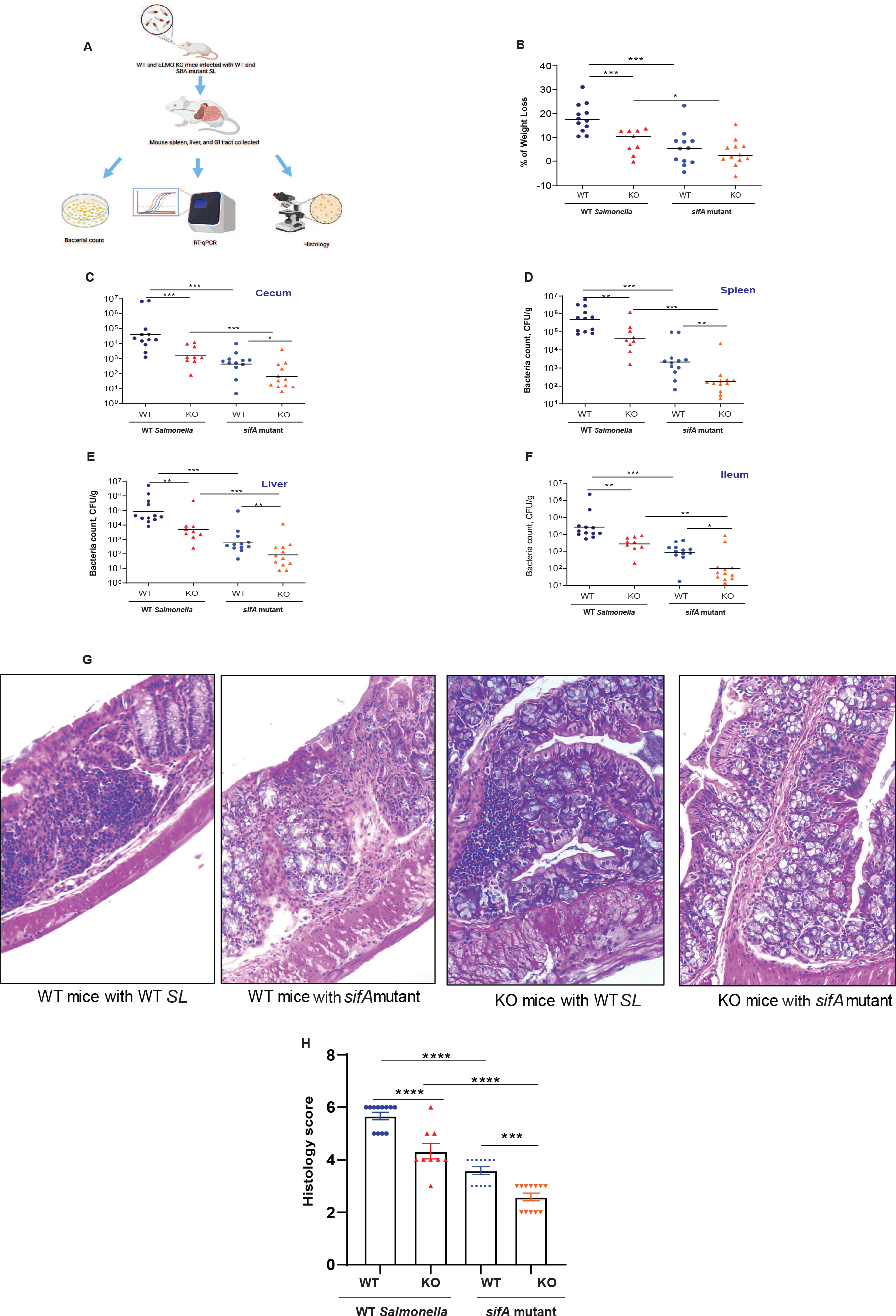
Infection of WT and global ELMO1KO mice with WT *Salmonella* (*SL*) and *sifA* mutant shows the involvement of ELMO1-SifA interaction in bacterial dissemination and inflammatory responses. (A) Schematic diagram represents the experimental design. (B) The percentage of weight loss was measured in WT and global ELMO1 KO mice infected via oral gavage with WT *SL* and *sifA* mutant strains for 5 days. (C-F) Bacterial burden was assessed at day 5 of infection in the cecum (C), spleen (D), liver (E), and ileum (F) of WT and global ELMO1 KO mice infected with WT *SL* and *sifA* mutant strains. (G-H) The H&E staining (G) and the pathology score (H) were assessed based on the degree of crypts loss, the infiltration of leukocytes in both mucosa and submucosa of WT and global ELMO1 KO mice infected with WT *SL* and *sifA* mutant. *, **, ***, and **** mean *p* ≤ 0.05, ≤ 0.01, ≤ 0.001, and ≤ 0.0001, respectively as assessed by Mann Whitney test.

Five days after infection, the percentage of weight loss was significantly higher in WT mice compared to ELMO1 KO mice infected with WT SL, and higher weight loss was recorded in mice infected with WT SL compared to the littermates infected with the *sifA* mutant **(Figure 2B)**. The bacterial burden in the cecum, spleen, liver, and ileum **(Figure 2C-2F)** was lower in mice infected with the the *sifA* mutant compared to mice infected with WT *SL*, with a significant decrease in the bacterial load in ELMO1 KO mice compared with WT mice **(Figure 2C-2F)**. H&E staining demonstrated loss of crypts and dense infiltration of leukocytes in both mucosa and submucosa of the infected mice ileum with a higher inflammatory infiltrate in WT *SL*-infected mice compared to mice infected with the *sifA* mutant **(Figure 2G)**. The histology score was significantly higher in WT mice compared to ELMO1 KO mice regardless of the inoculated strain **(Figure 2H)**. Overall, the degree of infection and inflammation was lower in mice infected with the *sifA* mutant, and it was the lowest in ELMO1 KO mice infected with the *sifA* mutant.

Since ELMO1 is expressed both in epithelial cells and myeloid cells, we aimed to assess the effect of ELMO1 expression on myeloid cells, its interaction with SifA *in vivo*, and the impact of this interaction on *Salmonella* dissemination. To this end, we orally infected WT and ELMO1 KO in myeloid cell-specific (LysM-cre-driven) (Das et al., 2015) mice with WT *SL* or the *sifA* mutant for 5 days. Similar to the global KO mice, the bacterial burden in the spleen and ileum was lower in mice infected with the *sifA* mutant compared to mice infected with WT *SL*, with a significant decrease in the bacterial load in LysM-cre-ELMO1 KO mice compared with WT mice **(Supple Figure 2)**. Next, we assessed the relevance of ELMO1 and SifA interaction in the early phase of infection (infection by gavage for 2 days) in WT and global ELMO1 KO mice. In WT mice, WT *SL* infection caused a significant weight loss and higher bacterial burden in the cecum, liver, spleen, but not in the ileum, compared to the *sifA* mutant **(Supple Figure 3 A-E)**. In contrast, we could not find any difference between WT *SL* and the *sifA* mutant in ELMO1 KO mice **(Supple Figure 3 A-E)**. The bacterial burden and weight loss were slightly higher in WT mice compared to ELMO1 KO mice in WT *SL* infection **(Supple Figure 3 A-E)**.

To assess the role of ELMO1-SifA interaction in regulating inflammatory responses and in the induction of innate immune responses *in vivo*, the expression of inflammatory transcripts such as TNF-α, MCP-1, IL-6, IL-1β, and CXCL-1 was assessed by RT-qPCR in the cecum **(Figure 3 A-E**) and spleen **(Figure 3 F-J)** of WT mice and ELMO1 KO mice infected with WT *SL* or the *sifA* mutant. The expression of inflammatory cytokines was significantly reduced in mice infected with the *sifA* mutant compared to WT *SL* infected mice, with much decrease in the levels of these transcripts in ELMO1 KO mice compared with the WT mice in both infections. Similar to the bacterial burden and histology score, the expression of inflammatory cytokines had the similar finding where WT mice infected with WT *SL* showed the highest pro-inflammatory cytokines and the ELMO1 KO mice infected with *sifA* mutant had the lowest pro-inflammatory cytokines. **(Figure 3)**. In early kinetic of infection (after 2 days of infection), the transcript level of inflammatory cytokines was higher in WT mice infected with WT *SL* compared to the other groups. ELMO1 KO mice infected with the *sifA* mutant showed the lowest level of inflammatory transcripts **(Supple Fig 4)**.

**Figure 3:**
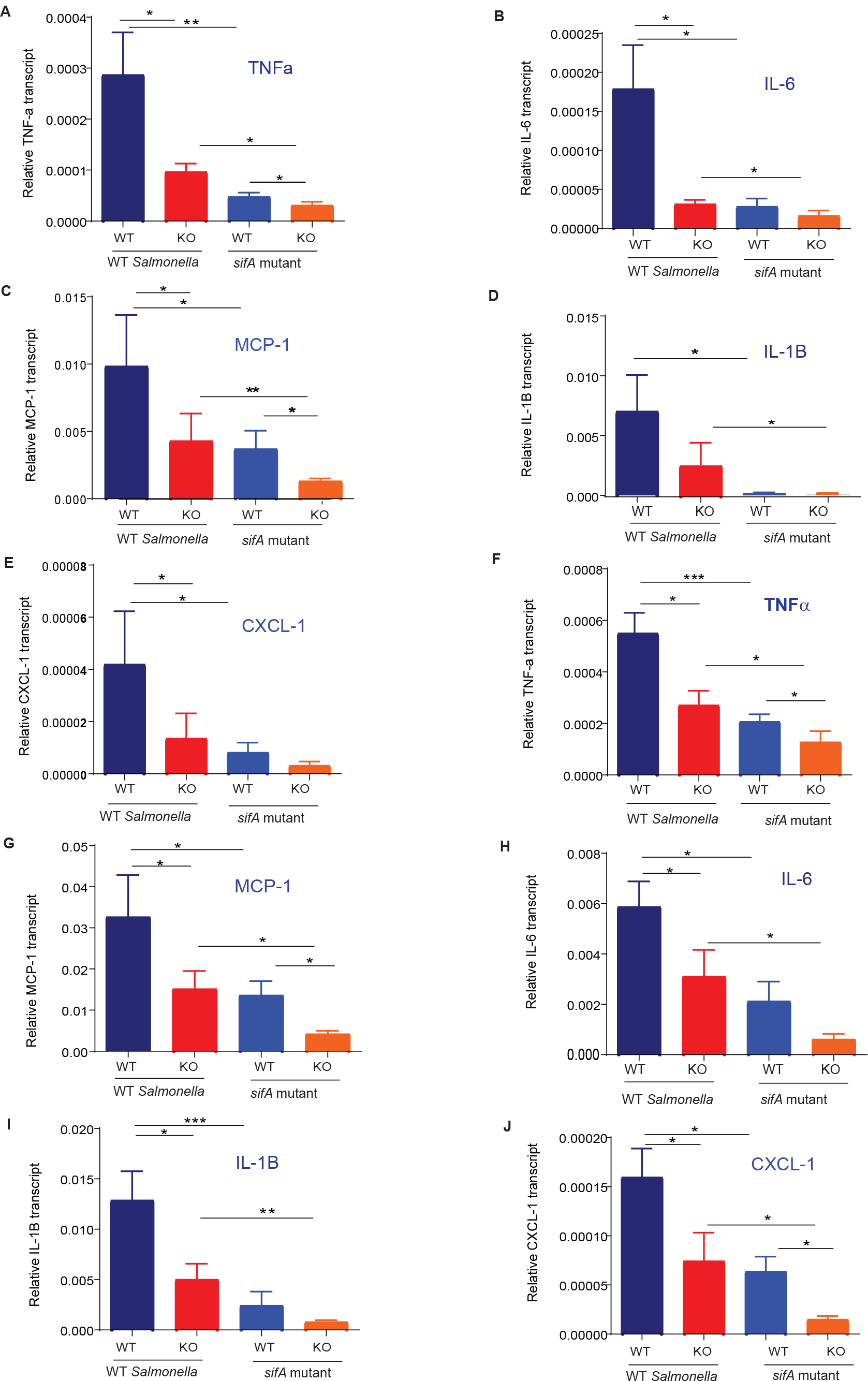
Infection of WT and global ELMO1 KO mice with WT *SL* and *sifA* mutant shows the involvement of ELMO1-SifA interaction on the induction of innate immune responses. (A-E) Total RNA was isolated from the cecum of WT and global ELMO1 KO mice infected with WT *SL* and *sifA* mutant for 5 days as in Figure (2), the transcript level of inflammatory cytokines such as TNF-α (A), IL-6 (B), MCP-1 (C), IL-1β (D), and CXCLl-1 (E) was measured by RT-qPCR. (F-J) Total RNA was isolated from the spleen of WT and global ELMO1 KO mice infected with WT *SL* and *sifA* mutant in Figure (2), the transcript level of inflammatory cytokines such as TNF-α (F), MCP-1 (G), IL-6 (H), IL-1β (I), and CXCLl-1 (J) was measured by RT-qPCR. *, **, *** means *p* ≤ 0.05, ≤ 0.01, and 0.001, respectively as assessed by Mann Whitney test.

**Figure 4.**
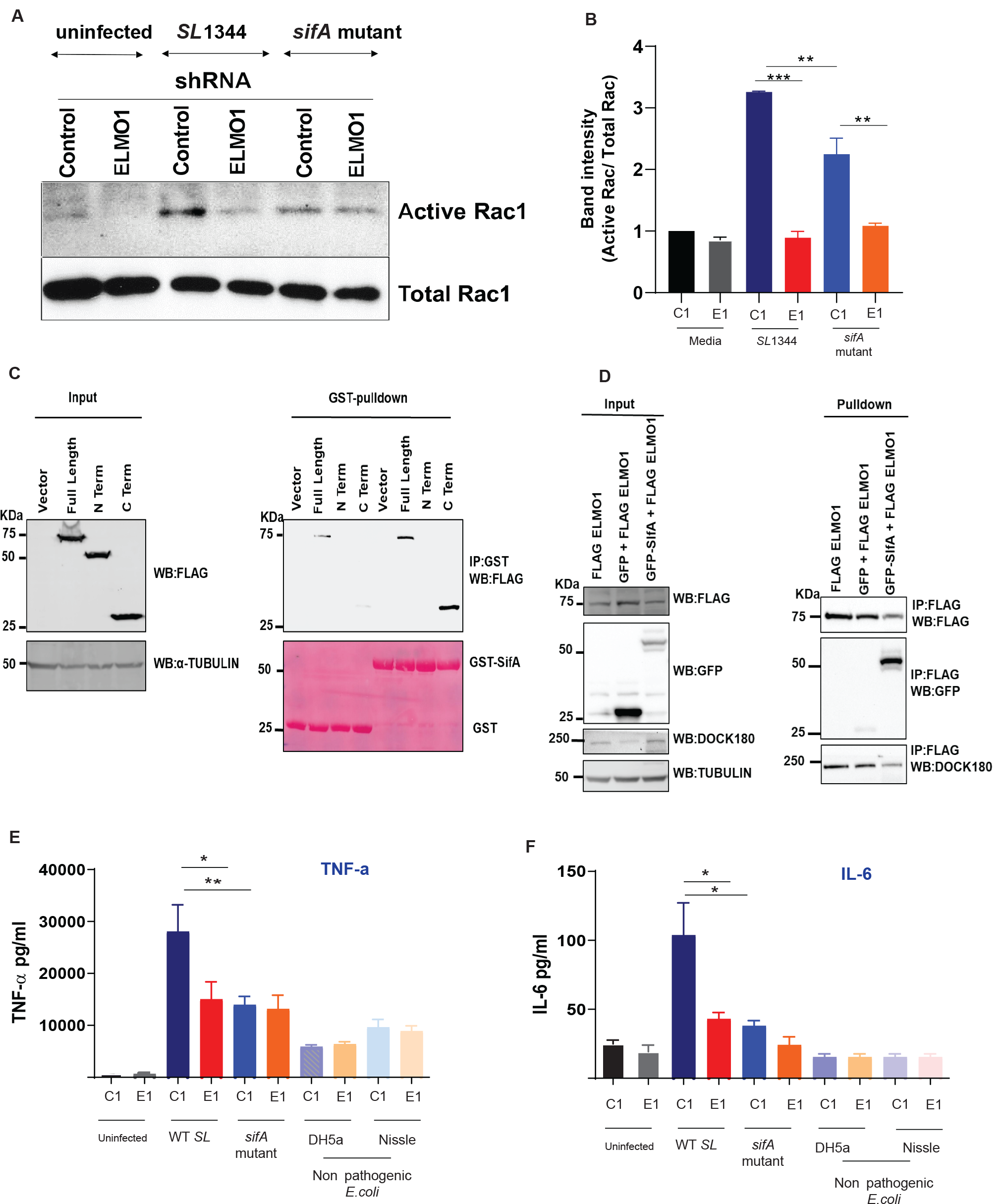
Effect of ELMO1-SifA interaction on induction of immune responses in control and ELMO1 shRNA cells. (A) Control (C1) or ELMO1 shRNA (E1) J774 cells were infected with WT *SL* or *sifA* mutant strain for 1 h, Rac1 Activity was assessed by pull down assay using GST-PBD beads, followed by WB for active (GTP bound) Rac1 (upper panel) and total Rac1 (lower panel) using Rac1 Antibody. (B) The band intensity for active Rac1 in was normalized to total Rac1 and the ratio of active Rac1/ total Rac1 was quantified in all samples. The densitometry was done with 3 individual experiments **. *** means *p*< 0.01 and 0.001, respectively. (C) HEK293 cells transfected with FLAG tagged ELMO1-full length (FL), FLAG tagged ELMO1-N terminal (NT) (aa-1-532) and FLAG tagged ELMO-1-C terminal (CT) (aa-532-727) of ELMO1, then the cell lysates incubated with GST-SifA beads. (left) represents pulldown assays using anti-Flag antibody (upper panel) and Ponceau S stain (lower panel). (Right): Western blot of the input cell lysates using anti-Flag antibody (upper panel), and α-Tubulin was used as a loading control (lower panel). (D) Co-immunoprecipitation assay to check any interference in the binding of Dock180 when ELMO1 interacts with SifA. FLAG-ELMO1 and GFP-SifA were transfected in HEK293 cells followed by cell lysis and IP using anti-FLAG antibody. Proteins were visualized by immunoblotting with corresponding antibodies. (E-F) Control (C1) or ELMO1 shRNA (E1) J774 cells were infected with WT SL1344, *sifA* mutant strain, and non-pathogenic *E*.*coli* for 3 h, then the level of inflammatory cytokines such as TNF-α (E), IL-6 (F), was measured by ELISA in the supernatant of infected cells. *, ** means *p* ≤ 0.05, and ≤ 0.01, respectively as assessed by one-way ANOVA multiple comparisons.

### Impact of ELMO1-SifA interaction on the macrophage immune response *in vitro*

ELMO1 binds and stabilizes Dock180, which in turn activates Rac1 (Lu et al., 2004). Interestingly, we found that ELMO1 activates Rac1 during *Salmonella* infection (Das et al., 2015). To assess the impact of ELMO1-SifA interaction on Rac1 activation, control and ELMO1 shRNA J774 cells were infected with WT *SL* or the *sifA* mutant strain for 1 h **(Figure 4A)**. The amount of active Rac1 was higher after infection with WT *SL* in control shRNA cells compared to ELMO1 shRNA cells. The *sifA* mutant showed a reduction of Rac1 activity compared to the WT *SL* strain. The densitometry in **Figure 4B** confirmed that ELMO1 shRNA cells have the lowest active Rac1, and the level was comparable after infection with WT *SL* or the *sifA* mutant. These results suggest that ELMO1 may interact with other effectors and controls the active Rac1.

Since the N-terminal part of Dock180 interacts with the C-terminal part of ELMO1 (Grimsley et al., 2006; Komander et al., 2008) and controls Rac1 activity, we investigated the interaction of SifA with ELMO1 using the N-terminal and C-terminal part of ELMO1. GST pull-down with GST-SifA showed that SifA is bound by both full-length ELMO1 and the C-terminal part of ELMO1, but not the N-terminal part of ELMO1 (**Figure 4C)**. As Dock180 and SifA both bind to ELMO1 in the C-terminus, we checked whether there is any interference between their interaction. The co-immunoprecipitation of FLAG-ELMO1 and GFP-SifA in HEK293 cells showed that Dock180 can interact with ELMO1 in the presence of SifA **(Figure 4D)**.

Next, we assessed the impact of ELMO1-SifA interaction on the immune response generated from the macrophages *in vitro*. We measured the level of inflammatory cytokines such as TNF-α and IL-6 in control and ELMO1 shRNA J774 cells after infection with WT *Salmonella*, with the *sifA* mutant, or with non-pathogenic bacteria such as *E. coli* Nissle-probiotic strain and *E. coli* DH5α. As expected, we did not detect any difference in the level of cytokines in the control and ELMO1 depleted cells infected with non-pathogenic bacteria **(Figure 4E-4F)**. Similarly, the level of TNF-α and IL-6 was comparable in control and ELMO1 shRNA cells infected with the *sifA* mutant **(Figure 4E-4F)**. On the other hand, the level of TNF-α and IL-6 was significantly higher in control cells compared to ELMO1 shRNA cells during the infection with WT *SL* **(Figure 4E-4F)**, and the level of these cytokines was significantly higher in control cells infected with WT *SL* compared to control cells infected with the *sifA* mutant **(Figure 4E-4F)**.

### Other enteric bacterial effectors containing WxxxE motif interact with ELMO1 and controls pro-inflammatory cytokines

To assess if ELMO1 interacts with WxxxE motif-containing effectors from *Shigella* (IpgB1, IpgB2) and *E*.*coli* (Map), we performed a GST pull-down assay using GST-IpgB1, GST-IpgB2, and GST-MAP with purified His-ELMO1 full length (FL) **(Figure 5A)**. We found that ELMO1 interacts with all these bacterial effectors **(Figure 5A)**. Next, we assessed the effect of overexpression of these effectors in control (C1) and ELMO1 (E1) shRNA J774 cells upon stimulation with bacteria and/or bacterial products such as LPS. To this end, control and ELMO1 shRNA cells were transfected with GFP-tagged bacterial effectors (SifA, IpgB1, IpgB2, MAP). The overexpression of the bacterial effectors was confirmed by WB and it was comparable in control and ELMO1-depleted cells and the interaction of bacterial effectors with endogenous ELMO1 was specific as there was no expression of ELMO1 in ELMO1 shRNA cells **(Figure 5B)**. To assess the relevance of the ELMO1-bacterial effectors interaction on the immune response, we infected some of the transfected cells from **Figure 5B** with *E. coli* K12, a non-pathogenic bacterium that lacks the WxxxE motif, and we measured TNF-α levels in the supernatant. The level of TNF-α was increased with infection, and it was significantly higher in control (C1) cells transfected with SifA, IpGB1, IpGB2, and/or MAP compared to ELMO1 (E1) shRNA cells **(Figure 5C)**. In contrast, the level of TNF-α was comparable in control and ELMO1 shRNA cells transfected with GFP empty vector **(Figure 5C)**. Next, we stimulated with LPS both control and ELMO1 shRNA cells transfected with the bacterial effectors, and we measured TNF-α and IL-6 in the supernatant. Similarly, the level of TNF-α and IL-6 was increased following LPS challenge, and the level of these cytokines was significantly higher in control (C1) cells transfected with bacteria effectors compared to transfected ELMO1 (E1) shRNA cells **(Figure 5D-5E)**.

**Figure 5:**
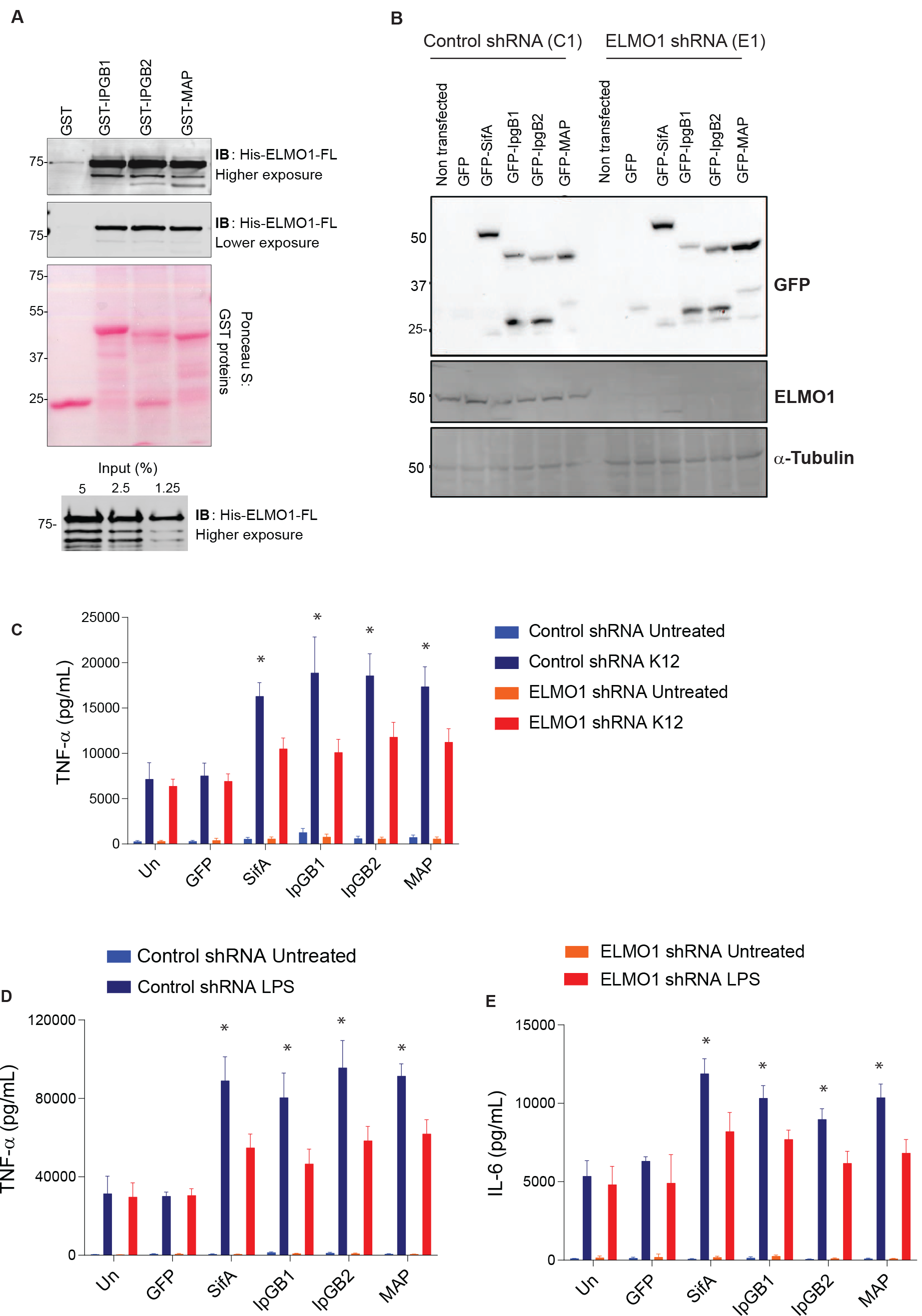
Interaction of ELMO1 and bacterial effectors with WxxxE signature motif controls inflammatory responses. (A) (upper panel) Recombinant His-ELMO1-full length (FL) was used in a GST pulldown assay with GST, GST-IpgB1, GST-IpgB2, or GST-Map. The bound ELMO1-FL was visualized by immunoblot (upper panel). Equal loading of GST proteins was confirmed by Ponceau S staining (lower panel). (lower panel) The input of His ELMO-1-FL was shown. (B) Control (C1) or ELMO1 (E1) shRNA J774 cells were transfected with GFP vector control, GFP-SifA, GFP-IpgB1, GFP-IpgB2, and GFP-Map. The cell lysates were assessed by anti-GFP antibody (upper panel), and anti-ELMO1 antibody (middle panel). Equal loading was confirmed by α-Tubulin (lower panel). (C-E) Control or ELMO1 shRNA J774 cells were transfected with GFP vector, GFP-SifA, GFP-IpgB1, GFP -IpgB2, and GFP-Map as in (B), and then the cells were challenged with *E*.*coli* K12 (C) or LPS (D-E). The supernatants were collected and the level of TNF-α (C, D) and IL-6 (E) was measured by ELISA. Data in C-E represent mean ± SEM of three independent experiments. * means p ≤ 0.05 as assessed by one-way ANOVA multiple comparisons.

## DISCUSSION

The recent development of omics technology and the progress in the field of microbiology and cell biology established the importance of physiological homeostasis in the intestinal mucosa. To maintain homeostasis, host immune signaling pathways need to discriminate between commensal and pathogens that cause infections. So far, we know partly about bacterial evasion mechanism; how bacterial pathogens have evolved to avoid host immune defenses; and how pathogenic effectors downregulate inflammation to enable pathogens to reside within the host and avoid clearance (Alto and Orth, 2012; Brodsky and Medzhitov, 2008; Miao and Miller, 1999; Van Avondt et al., 2015). However, our knowledge is still limited on how host immunity discriminates between commensal and pathogenic microbes, balances the inflammatory cytokine responses, and helps the host to clear invading pathogens. Here, we have addressed how a host signaling pathway interacts with a subset of bacterial effectors, and how this interaction leads to inflammation, which may be important in controlling infections. Specifically, we have shown the interaction between the microbial sensor ELMO1 and a group of bacterial effectors that share a signature motif [WxxxE (Trp-x-x-x-Glu)]. Using *Salmonella* WxxxE effector SifA, we further investigated the impact of effector-host interactions in the ELMO1 KO mice model and in ELMO1-depleted macrophages.

Alto and colleagues described a large family of 24 WxxxE effectors present in enteric pathogens that mimic GTPases and use the downstream signaling of RhoA, Rac1, and Cdc42 without the need for GTP (Alto et al., 2006). The bacterial effectors include SifA, SifB of *Salmonella*; IpgB1, IpgB2 of *Shigella*; Map, EspT, and EspM of *E. coli*, which share the unique WxxxE motif **(Figure 1)**. TheBLAST search revealed that the WxxxE motif is widely distributed among enteric, non-enteric and plant pathogens, but is absent in commensals. **(Figure 1, Supplementary Fig 1A-1D)**. Interestingly, the TIR domain of human TLR proteins has also the WxxxE motif. Two type III effector proteins in plant pathogens, WtsE and AvrE, require the WxxxE motif for disturbing host pathways by mimicking activated host G-proteins (Ham et al., 2009). In enteric pathogens, WxxxE effectors functionally mimic the Rho family GTPase (Alto et al., 2006; Stebbins and Galán, 2001) and effectors such as IpgB1, IpgB2, and Map are mostly involved in the rearrangements of host cytoskeletal processes, probably to facilitate bacterial entry (Alto et al., 2006; Handa et al., 2007). The *Brucella* effectors BtpA or BtpB with the TIR domain have the WxxxE motif and are involved in protection against microtubule depolymerization (Felix et al., 2014). It is not known whether all WxxxE effectors share a similar function or play different roles depending on the pathogen. Our work will provide new insight into this functional aspect of pathogenesis.

The *Salmonella* effector SifA is secreted by the type-III secretion system encoded on the *Salmonella* pathogenicity island 2 (SPI-2). It is essential for the formation of the *Salmonella*-containing vacuole (SCV) and for extending the Sif membrane networks and the survival of the bacteria inside macrophages (Beuzón et al., 2000; Ohlson et al., 2008; Stein et al., 1996). SifA has the WxxxE motif (Alto et al., 2006; Ohlson et al., 2008), and its C-terminus is essential for maintaining the tertiary structure of SifA, which is crucial for the interaction with the SKIP protein important for microtubule formation (Diacovich et al., 2009). Furthermore, the W197 and E201 residues of the WxxxE are required for the binding of SifA with SKIP leading to the antagonism of G-protein Rab9 (Jackson et al., 2008). We found that the WxxxE motif is also important for the interaction between SifA and ELMO1. Since ELMO1 interacts with bacterial effectors containing WxxxE, these effectors can modulate host immune signaling to produce the appropriate response. Our results reveal that ELMO1 plays a role in the internalization of both pathogenic and non-pathogenic bacteria but stimulates the release of pro-inflammatory cytokines (e.g., TNF-α) only in response to pathogenic bacteria. Collectively, these findings indicate that ELMO1-WxxxE effector interaction could explain the differential immune response mediated by ELMO1 to discriminate between pathogens and commensals.

ELMO1 facilitates intracellular bacterial sensing and the induction of inflammatory responses following enteric infection (Das et al., 2011; Das et al., 2015; Sarkar et al., 2017; Sayed et al., 2020). To gain additional insights, we evaluated the relevance of ELMO1-SifA interaction on bacterial internalization, pathogenesis, and host immune responses. Using WT and ELMO1 KO mice (global KO and myeloid cell specific ELMO1 KO) challenged with WT *Salmonella* or the *sifA* mutant, we found that the infection was less severe in ELMO1 KO mice, particularly in mice infected with the *sifA* mutant compared to WT *Salmonella*. Our findings suggest that ELMO1-SifA interaction increases bacterial colonization, dissemination, histopathology score, and inflammatory response *in vivo*. It is possible that the caspase-3 cleavage of SifA may be required for bacterial dissemination (Patel et al., 2019), and that there may be a link with ELMO1, but this needs to be further investigated. Previous work from us showed that ELMO1 is required for maximal bacterial internalization and pro-inflammatory responses during enteric infection (Das et al., 2015). Here, we have further dissected the function of ELMO1 and showed the importance of specific bacterial effectors. It is already known that deletion of SifA impairs the ability of *Salmonella* to invade and replicate within the host cells, evade the host immune system, and disseminate to extraintestinal organs. Our study also correlates with the results from Patel *et al*. (Patel et al., 2019) where they have reported that deletion of the SifA from *Salmonella* results in a 1-log lower dissemination of *Salmonella* to the liver. Therefore, the degree of infection and inflammation was decreased in mice infected with the *sifA* mutant compared to mice infected with WT *Salmonella*. Since ELMO1 is crucial to bacterial internalization and the induction of inflammatory response, infection with WT *Salmonella* was more attenuated in ELMO1 KO mice. ELMO1-depleted phagocytes exhibited significantly lower bacterial clearance, which also correlated to lower activity of lysosomal enzymes compared to control phagocytes (Sarkar et al., 2017). In addition, SifA stimulates host signaling analogous to the Rab GTPases, which is crucial to regulate membrane trafficking to the intracellular vacuole housing *Salmonella* in the host cytoplasm (Beuzón et al., 2000; Boucrot et al., 2005; Ruiz-Albert et al., 2002). Future cell biology and structural biology studies are required to understand the detailed downstream effects of ELMO1-SifA interaction in the maintenance of *Salmonella* vacuole and any endo-lysosomal machineries that control bacterial clearance.

Our study has shown the interaction between ELMO1 and bacterial effectors containing the WxxxE motif, and how this interaction is important for the generation of inflammation. We evaluated the impact of the interaction between ELMO1 and the WxxxE containing bacterial effectors on the inflammatory response released from macrophages *in vitro*. We found that the production of pro-inflammatory cytokines such as TNF-α and IL-6 was lowered in ELMO1-depleted macrophages with significantly lower amount in cells infected with the *sifA* mutant compared to cells infected with WT *Salmonella*. Likewise, lower cytokine levels were released from ELMO1-depleted macrophage transfected with other WxxxE containing effectors such as IpgB1, IpgB2, and Map upon stimulation with bacteria or bacterial products (LPS). The level of inflammatory transcripts was low in ELMO1 KO mice and in mice infected with the the *sifA* mutant. In a parallel line, previous reports showed that the *sifA* mutant cannot replicate and survive in macrophages and, therefore, cannot stimulate an inflammatory response (Brumell et al., 2001; Knuff-Janzen et al., 2020). The common function of bacterial effectors is to down-regulate the immune signaling to hide inside the host. Here we have shown how ELMO1 interacts with SifA and other WxxxE effectors and activates inflammatory pathways to alert the host about pathogens. Prior studies have shown that the EPEC effectors EspT and Map activate NFκB and MAP kinase pathways to activate the immune signaling (Ramachandran et al., 2020; Raymond et al., 2011). We have shown that ELMO1 activates NFκB and MAP kinase pathways following *Salmonella* infections (Das et al., 2015). It will be important to understand the ELMO1-WxxxE effectors interaction that controls NFκB/MAPK-mediated inflammatory signals to trigger the production of cytokines.

In conclusion, our results in *Salmonella* have shown that ELMO1-SifA interaction promotes bacterial colonization, dissemination, and inflammatory immune response. We have provided evidence that ELMO1 can discriminate between enteric pathogens and commensal probably through interaction with the WxxxE containing effectors. The interaction between host engulfment protein ELMO1 and bacterial effectors is crucial to control the disease pathogenesis.

## MATERIALS AND METHODS

All methods involving animal subjects were performed in accordance with the relevant guidelines and regulations of the University of California San Diego and the NIH research guidelines.

### Bacteria and bacterial culture

*Salmonella enterica* serovar Typhimurium strain SL1344, *E. coli* strain K12 were obtained from the American Type Culture Collection (ATCC) (Manassas, VA, USA), *E. coli* DH5α was obtained from ThermoFischer Scientific, *E. coli* strain Nissle 1917 was obtained from Ardeypharm, and the SL1344 *sifA* mutant strain was obtained from Dr. Olivia Steele-Mortimer (Steele-Mortimer et al., 2002; Stein et al., 1996) . All bacteria were maintained as described previously (Das et al., 2015; den Hartog et al., 2021). Briefly, a single colony was inoculated into LB broth and grown for 8 h under aerobic conditions and then under oxygen-limiting conditions. For the SL1344 *sifA* mutant, streptomycin with a final conc of 100 μg/mL was added to LB broth. Cells were infected with a multiplicity of infection (moi) of 10.

### Cell culture and transfection

HEK293 cells and murine macrophage cell line J774 were obtained from American Type Culture Collection, (Manassas, VA, USA). Cells were maintained in high glucose DMEM (Life Technologies) containing 10% fetal bovine serum and 100 U/ml penicillin and streptomycin at 37°C in a 5% CO_2_ incubator. Control (C1) and ELMO1 depleted (E1) shRNA J774 cells were generated as previously described (Das et al., 2015) and maintained in complete media supplemented with 0.5 μg/ml Puromycin (Sigma). Cells were sub-cultured 24 hours prior to transfection. Transfections of plasmids were performed using Lipofectamine 2000 (Invitrogen) according to manufacturer’s protocol.

### Blast search and sequence alignment

Literature was searched to find published research regarding WxxxE motifs in plants, enteric, and non-enteric pathogens. Protein and bioinformatics databases UniProt (https://www.uniprot.org/), PATRIC (https://www.patricbrc.org/), and NCBI’s BLAST (https://blast.ncbi.nlm.nih.gov/Blast.cgi) were used to search for protein sequences and BLAST for related proteins that may have the WxxxE motif. Clustal Omega (https://www.ebi.ac.uk/Tools/msa/clustalo/) was used to align sequences and Adobe InDesign CS6 was used to design the final figures that combined all effectors from each category (enteric pathogens, non-enteric pathogens, plant pathogens, and human Toll-Like receptors).

### Infection of WT and ELMO1 KO mice with *Salmonella* strains

To assess the role of ELMO1 and of SifA effector protein on bacterial pathogenesis *in vivo*, age-and sex-matched WT and ELMO1 KO (either global KO, or LysM-cre driven where the ELMO KO was specifically deleted in myeloid cells) C57BL/6 mice were infected with *Salmonella enterica* serovar Typhimurium SL1344 wild-type or an isogenic *sifA* mutant (5×10^7^ cfu/ mouse) by oral gavage. The infection period was 5 days, and the mice were monitored for weight change and clinical signs of disease during this period. At day 5 post infection, the mice were sacrificed, and their tissues (liver, spleen, ileum, and cecum) were collected to assess bacterial colonization, inflammatory responses, and histology score. To assess early kinetic of infection, WT and ELMO1 KO were challenged with SL1344 or the *sifA* mutant by oral gavage and the infection lasted for 2 days. Animals were bred, housed, used for all the experiments, and euthanized according to the University of California San Diego Institutional Animal Care and Use Committee (IACUC) policies under the animal protocol number S18086. All methods were carried out in accordance with relevant guidelines and regulations and the experimental protocols were approved by institutional policies and reviewed by the licensing committee.

### Assessment of bacterial load and inflammatory response in the mice tissues

Harvested tissues were weighed first and then resuspended in phosphate-buffered saline (PBS), homogenized, diluted serially, and plated in LB agar plates. The *sifA* mutant was plated in LB agar plates with Streptomycin (Sigma, final conc 100 μg/mL). The bacterial load was calculated as colony forming unit (cfu) per gram of tissue. To assess the expression of inflammatory cytokines, tissue specimens from spleen, ileum, cecum, and liver were collected for RNA isolation from these tissues.

### RNA Preparation, Real-Time Reverse-Transcription Polymerase Chain Reaction

Total RNA was extracted using Direct-zol RNA MiniPrep Kit (Zymo Research, USA) according to the manufacturer’s instructions. cDNA was synthesized using the qScript™ cDNA SuperMix (Quantabio). Quantitative PCR (qPCR) was carried out using 2x SYBR Green qPCR Master Mix (Biotool™, USA) for target genes that normalized to the endogenous control gene using the 2^-ΔΔCt^ method. Primers were designed using NCBI Primer Blast software the Roche Universal Probe Library Assay Design software **(Table 1)**.

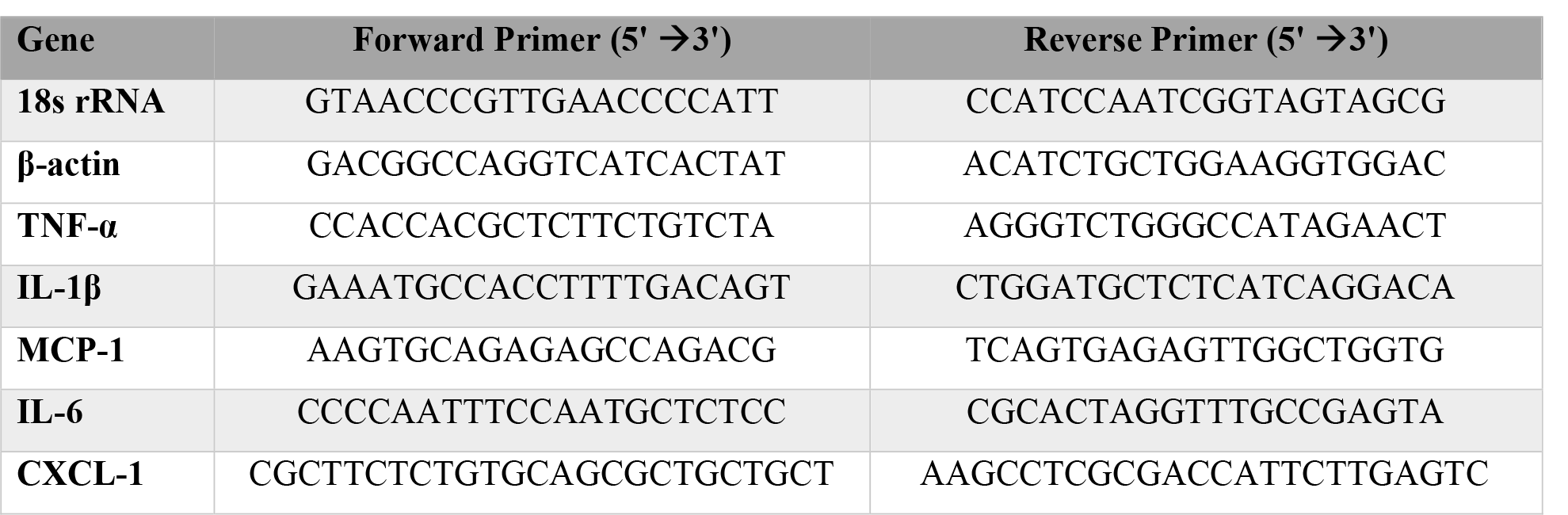

### Histopathology of WT and ELMO-1 KO mice following *Salmonella* infection

Ileal tissues from SL1344 and SL1344 *sifA*-infected WT and ELMO-1 KO mice were fixed using Zinc formalin and stained with H&E. The slides were evaluated for the presence of inflammatory cells such as neutrophils, mononuclear infiltrates, and mucosa architecture architecture, and the histology score for each slide was determined as described previously (Das et al., 2015; Sayed et al., 2020).

### Infection of J774 macrophages with *Salmonella* strains and nonpathogenic bacteria for cytokine assays

Control (C1) and ELMO1 (E1) depleted J774 macrophages cells were either left uninfected or infected with *Salmonella* strains (SL1344, and an isogenic *sifA* mutant), or nonpathogenic bacteria (*E. coli* strains K12, DH5α, and Nissle 1917). Supernatants were collected from the infected cells and tested for TNF-α, and IL-6, using a mouse ELISA kit (R&D Systems, USA) according to the manufacturer’s instructions.

### The impact of bacterial effectors on cytokine responses in J774 macrophages

Control (C1) and ELMO1-depleted (E1) J774 macrophages cells were transfected with plasmids containing the bacterial effectors (SifA, IpgB1, IpgB2, and Map) as described in the previous section. Transfected cells were challenged with LPS (100 ng/ml) for 6 h, and *E. coli* K12 (moi 10) for 3 h. Supernatants were collected at the time points and assessed for cytokines by ELISA.

### Bacterial internalization by gentamicin protection assay

Approximately 2×10^5^ cells were were seeded into 24-well culture dishes 18 hours before the infection and infected with bacteria with moi 10 for 1 hours in antibiotic-free media as described previously (Das et al., 2011; Das et al., 2015).

### Protein expression and purification

Recombinant His and GST-tagged proteins were expressed in *Escherichia coli* strain BL21 (DE3) and purified as previously described (Garcia-Marcos et al., 2009; Ghosh et al., 2008). Briefly, proteins in bacterial culture were induced with IPTG (0.5mM) overnight incubation at 25°C. Bacteria culture was centrifuged and cell pellet was lysed in either GST lysis buffer (25 mM Tris-HCl (pH 7.4) 20 mM NaCl, 1 mM EDTA, 20% (v/v) glycerol, 1% (v/v) Triton X-100, protease inhibitor cocktail) or His lysis buffer (50 mM NaH_2_PO_4_ (pH 7.4), 300 mM NaCl, 10 mM imidazole, 1% (v/v) Triton-X-100, protease inhibitor cocktail). Cell lysates were briefly sonicated and then centrifuged at 4°C for 30 mins at 12,000xg. Cleared cell lysates were then affinity purified using either glutathione-Sepharose 4B beads or HisPur Cobalt Resin, followed by elution and overnight dialysis in PBS. Proteins were then quantified and stored at -80°C.

### *In vitro* GST pulldown and immunoprecipitation

Recombinant purified GST-tagged proteins were immobilized onto the glutathione-Sepharose beads in binding buffer (50 mM Tris-HCl (pH 7.4), 100 mM NaCl, 0.4% (v/v) Nonidet P-40, 10 mM MgCl2, 5 mM EDTA, 2 mM DTT) for 2 hrs at 4°C with gentle agitation. GST-tagged protein bound beads were washed and incubated with purified His-tagged proteins or cell lysate in binding buffer overnight at 4°C with gentle agitation. The GST beads were then washed four times with NETN buffer (0.5% NP40, 0.1 mM EDTA, 20 mM Tris, pH 7.4, 300 mM NaCl) for cell lysates, or GST wash buffer (20 mM Tris, pH 7.4, 0.1 mM EDTA and 100 mM NaCl) for purified proteins.

For immunoprecipitation, transfected HEK293 cells were lysed in RIPA lysis buffer (50mM Tris pH 7.4, 150mM NaCl, 0.1% SDS, 1% NP40, 0.5% Sodium Deoxycholate) with 1X Proteinase Inhibitor Cocktail added immediately before use. Whole cell lysate was centrifuged to separate proteins from cell debris and quantified using the Lowry assay. One mg of total protein lysate was diluted 4 times its volume with cold 50mM Tris pH 7.4 (1X Proteinase Inhibitor Cocktail added fresh) and incubated with 40ul of antibody-conjugated beads overnight at 4°C with gentle agitation. Beads were washed 4 times with cold NP40 buffer (1% NP40, 0.5% Sodium Deoxycholate and 0.1% SDS in 1 X PBS) at 1500 rpm for 2mins.

Bound proteins were eluted by boiling beads for 10 mins at 95°C in 2X SDS-PAGE sample buffer containing β-mercaptoethanol. Proteins were separated using SDS-PAGE protein gel and transferred to Immobilon-P PVDF membrane. Proteins were visualized by immunoblotting with corresponding antibodies.

### Assessment of Rac1 activity in control and ELMO1 shRNA cells

The activity of Rac1 was assessed by using GST-PBD (glutathione S-transferase with p21-binding domain of Pak1) beads as described before (Das et al., 2011). Briefly, control and ELMO1 shRNA cells infected cells with WT SL or *sifA* mutant were lysed in buffer containing 50 mM Tris-HCl (pH 7.5), 2 mM MgCl2, 0.1 M NaCl, 1% NP-40, and 10% glycerol with protease inhibitors and incubated with GST coupled to PBD to precipitate Rac-GTP. The blots were visualized by electrochemiluminescence reagents (Pierce SuperSignal). The level of active Rac1 (GTP bound) was normalized to total Rac1 using ImageJ software.

### Statistical analysis

Results presented in this study were presented as the mean ± SEM and the bacteria load (CFU) was expressed as the geometric mean. Data compared using Mann–Whitney U test two-tailed Student’s t test and/or one-way ANOVA multiple comparisons as described in the specific positions. Results were analyzed in the Graph pad Prism and considered significant if p values were < 0.05.

## Acknowledgement

We are thankful to Dr. Neil Alto, UT Southwestern Dallas for providing the plasmids related to SifA, IpgB1, Map, Dr. Olivia Steele Mortimer, NIAID, Montana for providing the *SifA* mutant *Salmonella*, and Dr. Ulrich Sonnenborn from Ardeypharm for providing the *E. coli* strain Nissle 1917. We are grateful to Dr. Pradipta Ghosh, for providing suggestions during preparation of the manuscript. We are thankful to Dr. Gajanan Katkar, Dr. Stefania Tocci and Dr. Sajan Achi for reading the manuscript. We appreciate the technical support from Katherine Suarez, Eileen Lim, Mitchel Lau, Alicia Amamoto and Julian Tam, the lab interns of the Das lab.

## Data availability statements

Data available within the article or its supplementary materials.

## Supplementary Figures Legends

**Figure 1S:**
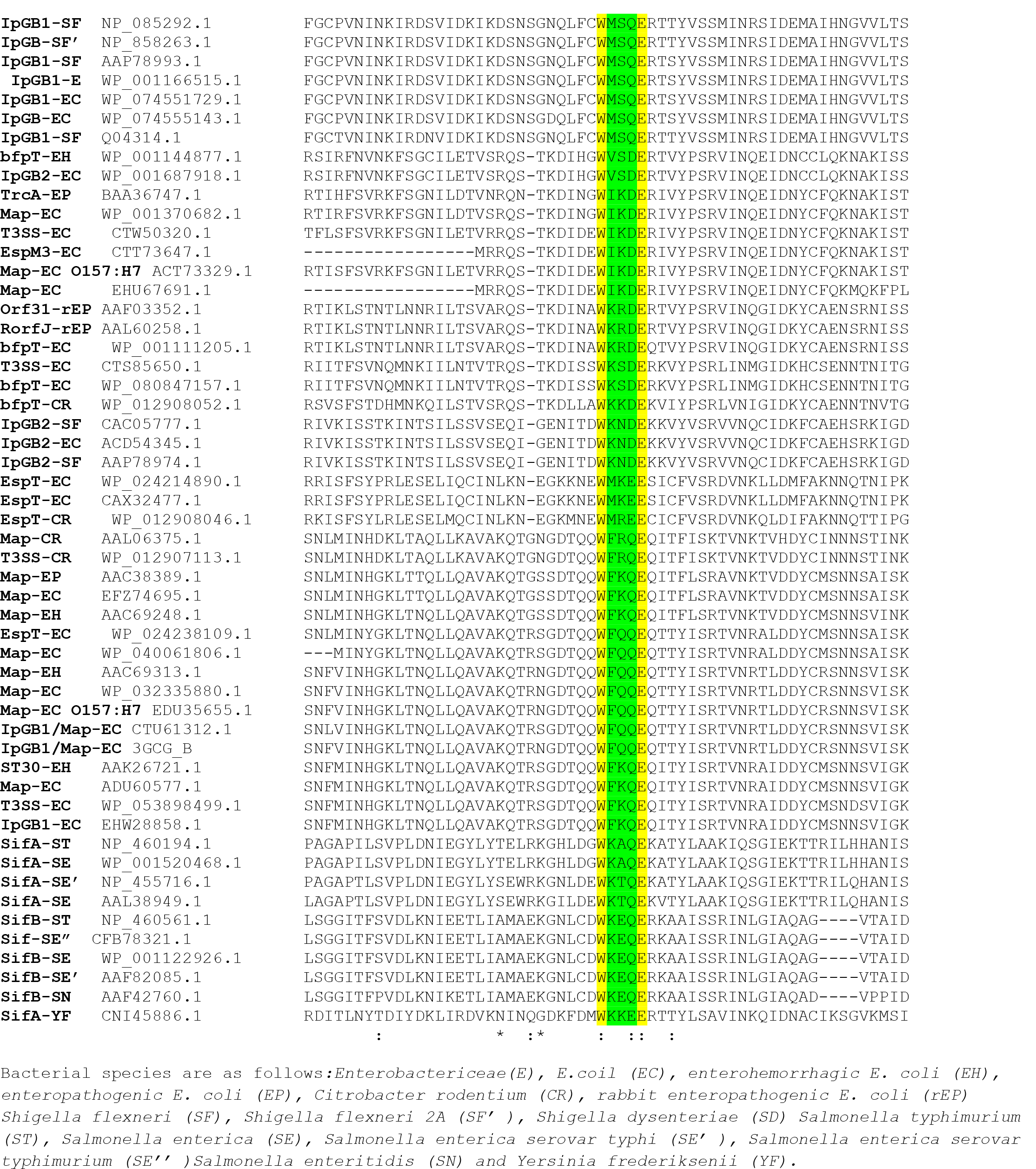
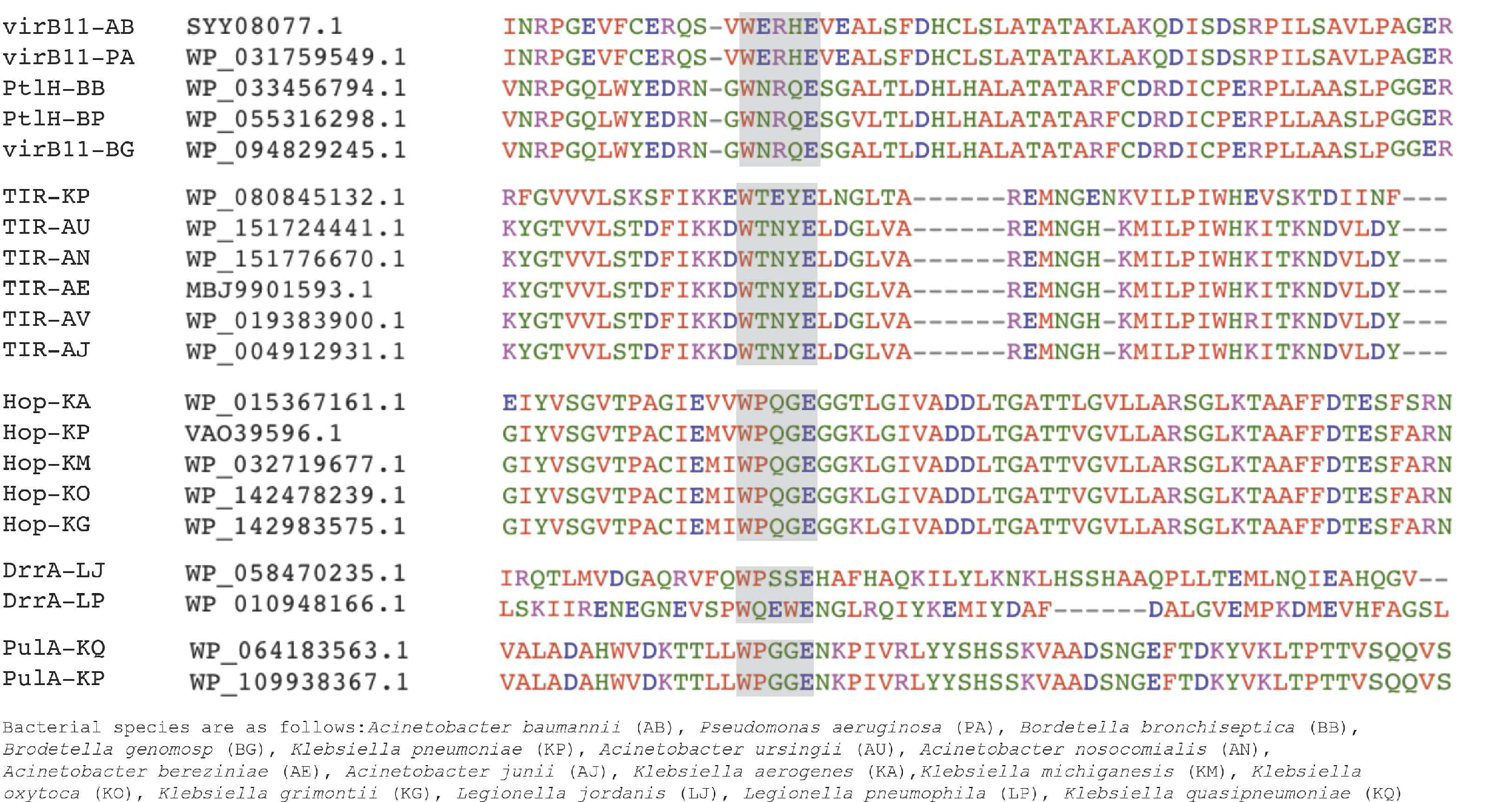
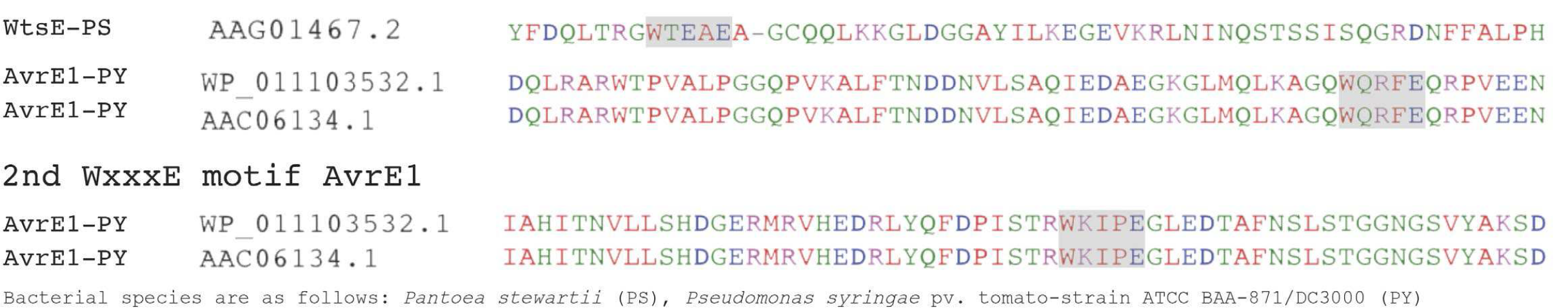
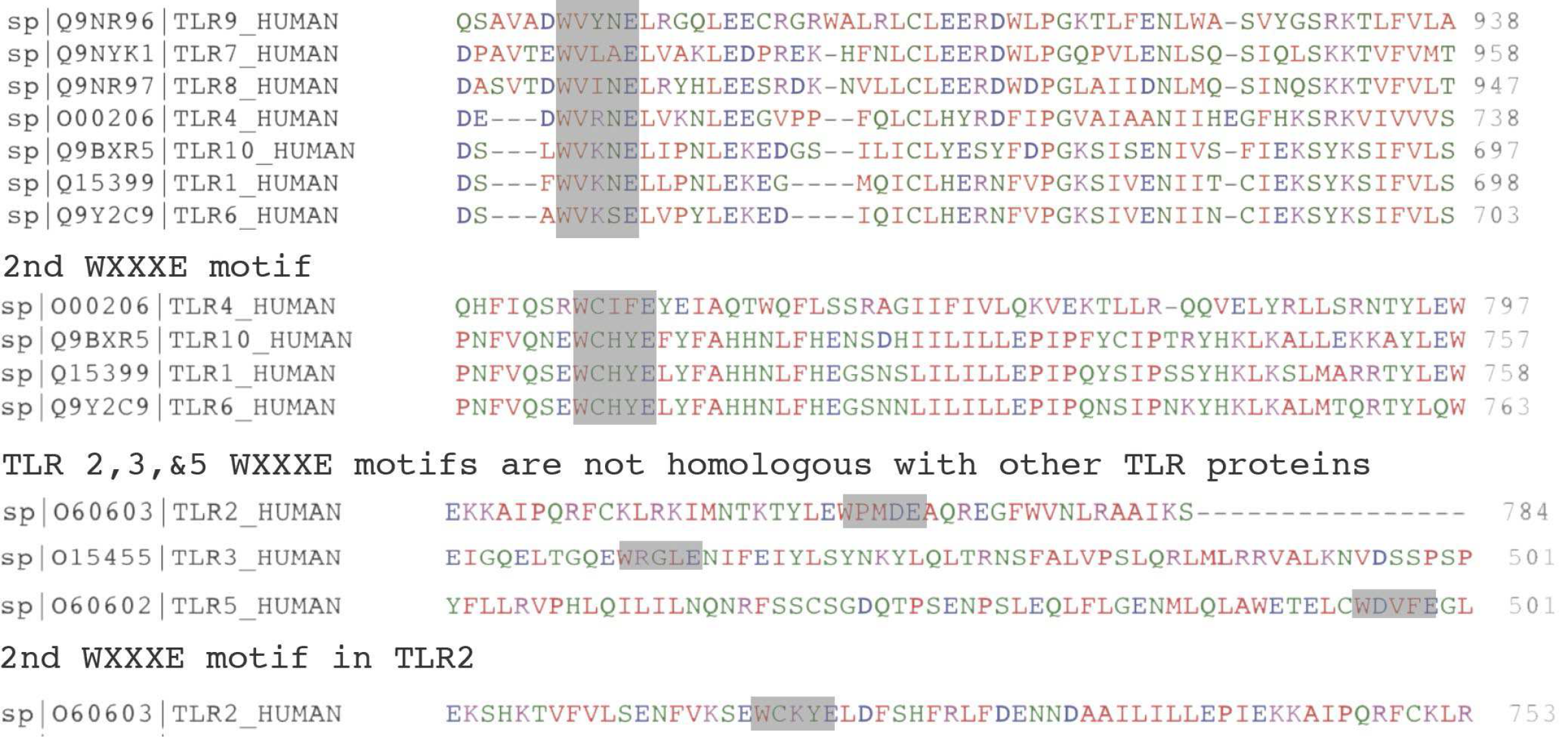
The distribution of the WxxxE motif among enteric, non-enteric, and host receptor. A BLAST search between amino acid sequences showed effectors with the WxxxE or “Trp-x-x-x-Glu” sequences within enteric pathogens (A), non-enteric pathogen (B), plant pathogens (C) and host proteins (D). (A) represents the list of enteric pathogens includes the name of effector and bacteria (left), the sequence ID (middle), and the amino acid sequences of the effectors (right). In the amino acid sequences WxxxE sequence is highlighted, where W and E are yellow highlighted, and xxx sequence is green highlighted. (B) represents the list of non-enteric pathogens includes the name of effector and bacteria (left), the sequence ID (middle), and the amino acid sequences of the effectors (right). (C) represents the list of plant pathogens includes the name of effector and bacteria (left), the sequence ID (middle), and the amino acid sequences of the effectors (right). (D) represents the list of human TLR containing this motif including the sequence ID and the name of receptor (left), and the amino acid sequences of the effectors (right).

**Figure 2S:**
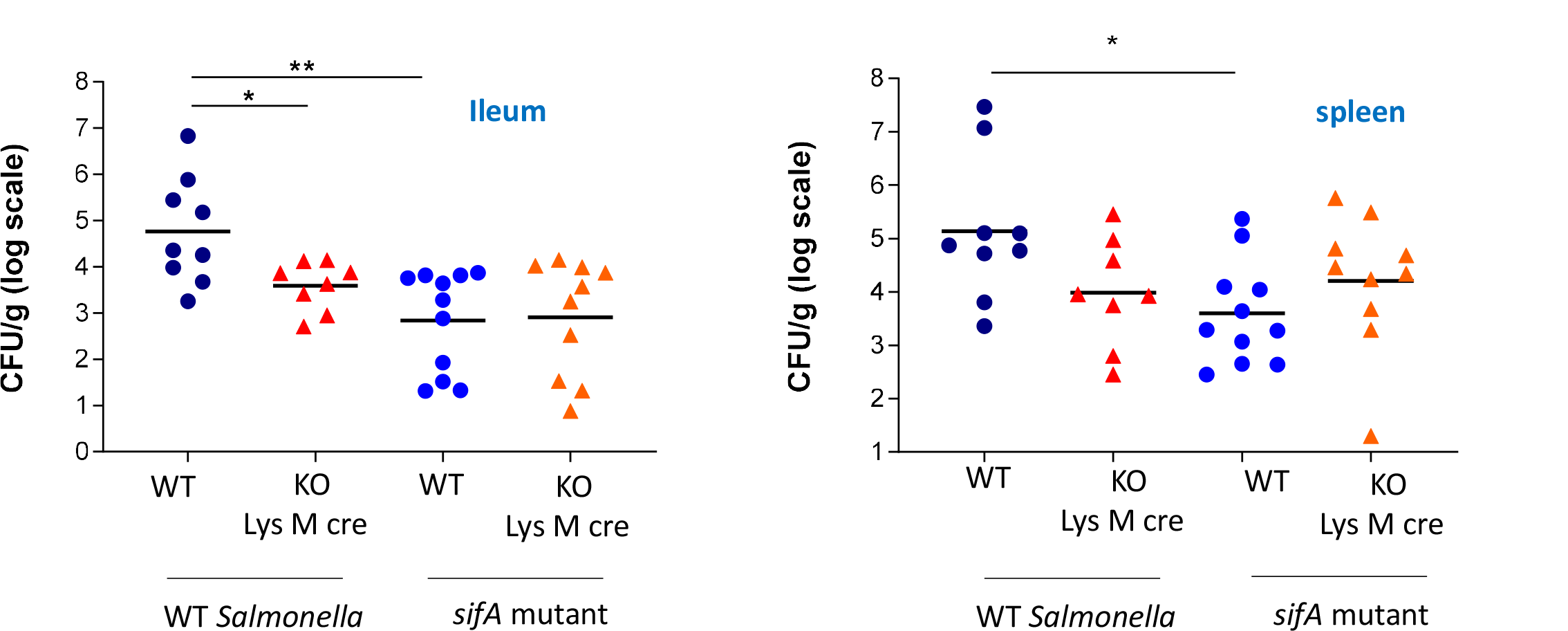
Effect of WT *SL* and *sifA* mutant infections on the bacterial colonization in Lys M-cre-driven ELMO1 KO mice. WT and myeloid cells specific ELMO1 KO mice (Lys M-cre) were infected via oral gavage with WT *SL* and *sifA* mutant strain for 5 days. (A-B) Bacterial burden was assessed at day 5 of infection in the ileum (A), and spleen (B) of the infected mice. *, ** mean *p* ≤ 0.05, and ≤ 0.01, respectively as assessed by Mann Whitney test.

**Figure 3S:**
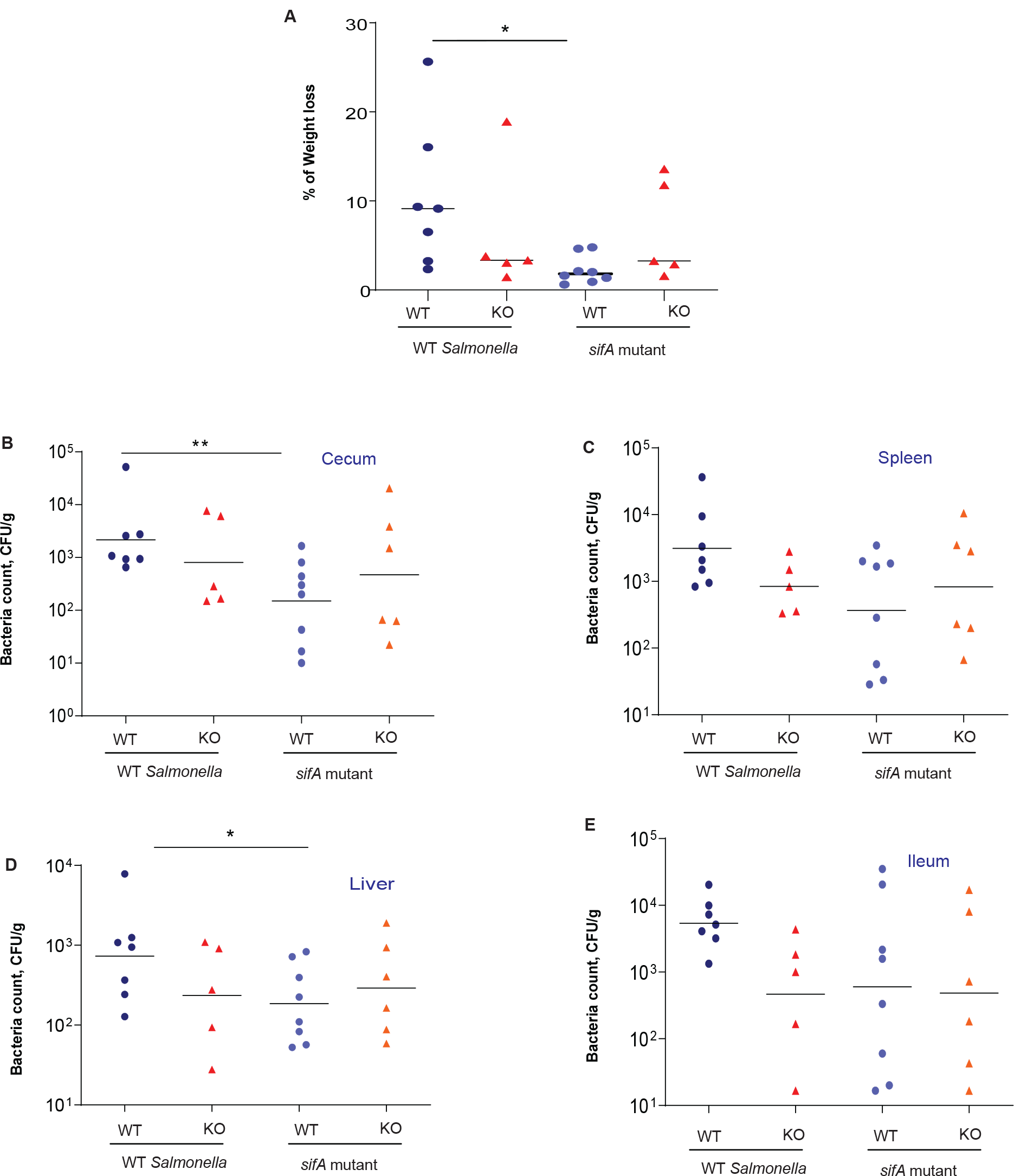
Effect of WT *SL* and *sifA* mutant infections for 2 days on the bacterial colonization in global ELMO1 KO mice. WT and global ELMO1 KO mice were infected via oral gavage with WT *SL* and *sifA* mutant strain for 2 days. (A) The percentage of weight loss was assessed in WT and global ELMO1 KO mice after 2 days of infection. (B-E) Bacterial burden was assessed at day 2 of infection in the cecum (B), spleen (C), Liver (D), and ileum (E) of the infected mice. *, ** mean *p* ≤ 0.05, and ≤ 0.01, respectively as assessed by Mann Whitney test.

**Figure 4S:**
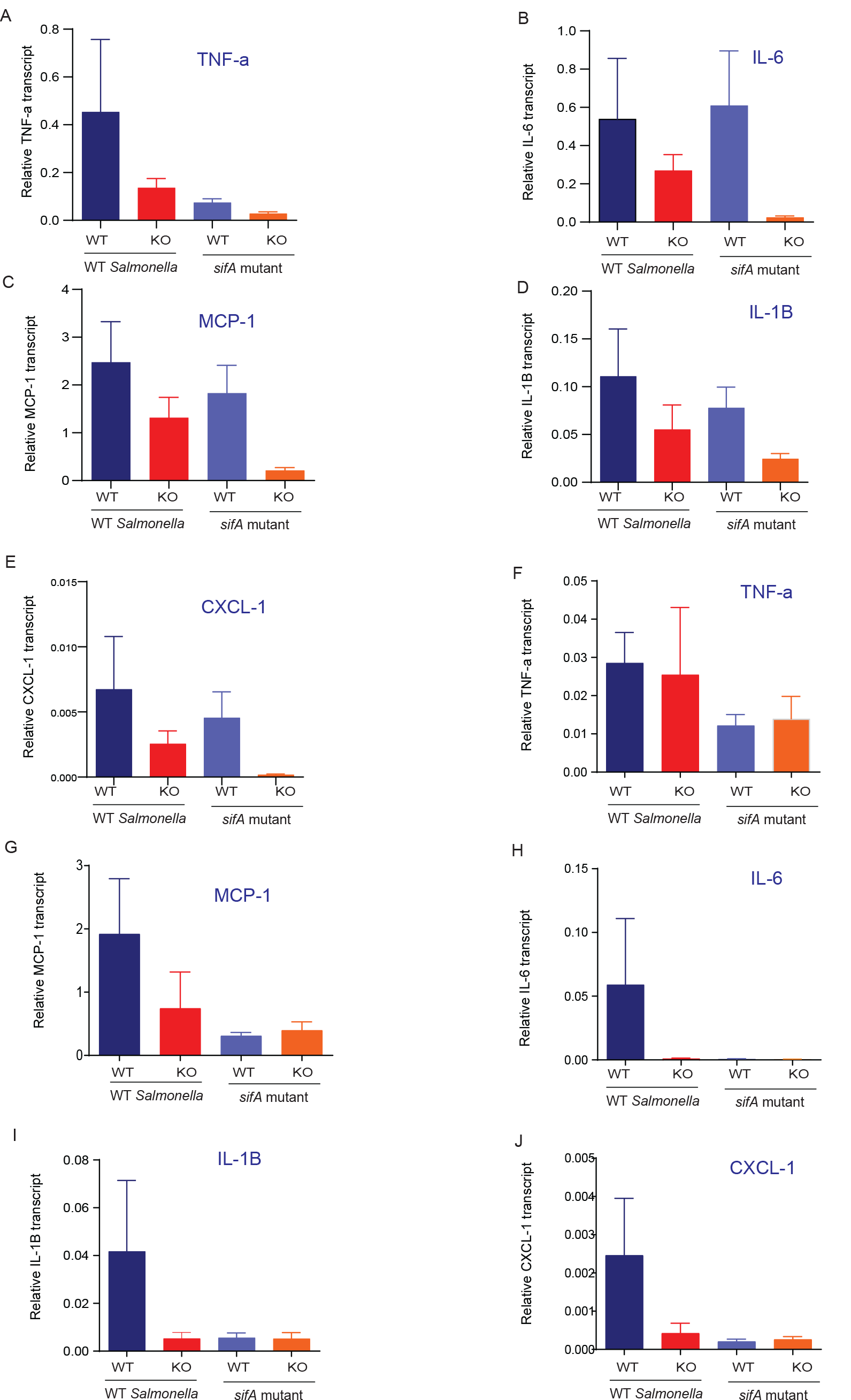
Effect of WT *SL* and *sifA* mutant infections for 2 days on the inflammatory cytokines in global ELMO1 KO mice. WT and global ELMO1 KO mice were infected via oral gavage with WT *SL* and *sifA* mutant strain for 2 days. Total RNA was isolated from the spleen and ileum of WT and global ELMO1 KO mice. (A-E) The transcript level of inflammatory cytokines: TNF-α (A), IL-6 (B), MCP-1 (C), IL-β (D), and CXCLl-1 (E) was measured by RT-qPCR in the spleen of infected mice. (F-J) The transcript level of inflammatory cytokines: TNF-α (F), MCP-1 (G), IL-6 (H), IL-1β (I), and CXCLl-1 (J) was assessed by RT-qPCR in the ileum of infected mice.

